# Infectious Disease Knowledge Graph (IDKG) for Antiviral and Antibacterial Drug Discovery

**DOI:** 10.64898/2026.11.12.766252

**Authors:** Holli-Joi Martin, Jon-Michael Beasley, Valeriia Kaneva, Ricardo Scheufen Tieghi, Kevin Causey, Alexander Tropsha, Eugene Muratov

## Abstract

Infectious diseases remain a leading cause of global morbidity and mortality, while therapeutic discovery is hindered by fragmented biomedical data and antimicrobial resistance. To address this challenge, we developed the Infectious Disease Knowledge Graph (IDKG), a queryable platform that integrates curated chemical, biological, pharmacological, and clinical information relevant to infectious-disease drug discovery. Using a customized ORION pipeline, we integrated 20 data sources, including DrugCentral, SMACC, Heli-SMACC, NPASS, ClinicalTrials.gov, and host–virus interaction databases, into a Neo4j graph containing more than 4.6 million nodes and 26 million edges. IDKG supports graph-based reasoning through Clinical Outcome Pathways (COPs) and incorporates curation workflows for small molecules, biologics, and complex mixtures. Illustrative applications include reconstruction of a literature-supported remdesivir pathway for COVID-19 and graph-based mechanistic interpretation of experimentally observed quorum-sensing inhibitor–antibiotic synergies in multidrug-resistant bacteria. The platform provides an interoperable framework for drug repurposing, target prioritization, and mechanism-based hypothesis generation. IDKG is freely available at https://idkg.mml.unc.edu.

## Introduction

Infectious diseases (IDs) remain a leading cause of morbidity and mortality worldwide^1^, presenting an urgent and evolving public health challenge driven by emerging pathogens, zoonotic spillovers, and the accelerating crisis of antimicrobial resistance^2,3^. Recent pandemics and the global spread of resistant microbes underscore the limitations of traditional drug discovery approaches, which rely heavily on target-based screening, take an average of 10–15 years, and cost over $2.2 billion per compound to bring new therapeutics to market^4,5^. The urgency to respond more rapidly is further hindered by the fragmented and siloed nature of biomedical data, limited interoperability across databases, and the labor-intensive nature of experimental validation^6^.

Concurrently, the explosion of heterogeneous biological data, ranging from genomics and transcriptomics to chemical biology, clinical trials, and real-world evidence, offers unprecedented opportunities for accelerating therapeutic discovery. However, its integration and actionable interpretation remain major bottlenecks. Knowledge graphs (KGs) have emerged as a powerful solution to these challenges^7–9^. By representing biomedical entities (e.g., drugs, targets, diseases, pathways) as interconnected nodes linked through semantically rich, evidence-based relationships, KGs enable machine-readable data integration, scalable hypothesis generation, and *in silico* exploration of complex biological systems^10^.

To address these needs in infectious-disease drug discovery, we developed the Infectious Disease Knowledge Graph (IDKG), a semantically integrated computational platform that consolidates chemical, biological, pharmacological, and clinical data relevant to infectious diseases. Building on ROBOKOP^8^, IDKG incorporates infectious-disease-focused resources including SMACC^11^, Heli-SMACC^12^, PathoPhenoDB^13^, and ClinicalTrials.gov^14^. The platform supports hypothesis-driven target identification, drug repurposing, mechanistic pathway exploration, and analysis of drug combinations. In this study, we describe the design and construction of IDKG and present illustrative antiviral and antibacterial use cases that illustrate how an infectious-disease-focused knowledge graph can support therapeutic-discovery workflows.

Relative to the general-purpose ROBOKOP framework, IDKG narrows the integration and curation strategy toward infectious-disease therapeutic discovery while retaining Biolink-compatible semantics. COVID-KOP extended ROBOKOP specifically for SARS-CoV-2 by adding COVID-19 literature and viral annotations,^18^ whereas KG-COVID-19 was developed as a configurable framework for constructing COVID-19-focused graphs for multiple downstream applications.^19^ IDKG differs in scope by integrating antiviral and antibacterial activity data, host–pathogen interactions, chemical structures and mixtures, clinical-trial information, and graph-based mechanistic reasoning within a single infectious-disease-focused resource.

## Results and Discussion

The Infectious Disease Knowledge Graph (IDKG) was developed to address the pressing need for integrated, high-resolution data infrastructure to support drug discovery for infectious diseases. Leveraging the foundational structure of ROBOKOP, IDKG was built using a modified version of the ORION^15^ pipeline (https://github.com/molecularmodelinglab/ORION_IDKG), enabling harmonization of diverse datasets into a semantically consistent, queryable graph. The current IDKG integrates curated data from 20 sources, including DrugCentral, SMACC, Heli-SMACC, NPASS, ClinicalTrials.gov, and HVIDB (**Table 1**). The resulting graph comprises over 4.6 million nodes and more than 26 million edges, representing complex interactions among drugs, targets, diseases, phenotypes, pathways, and clinical outcomes. A detailed breakdown of node types is provided in **Table 2**, with Organism Taxon (2,627,442), Small Molecule (1,382,871), and Protein (320,273) representing the largest categories. The predominant edge types connecting these node types are illustrated in Figure 1.

**Table 1.**
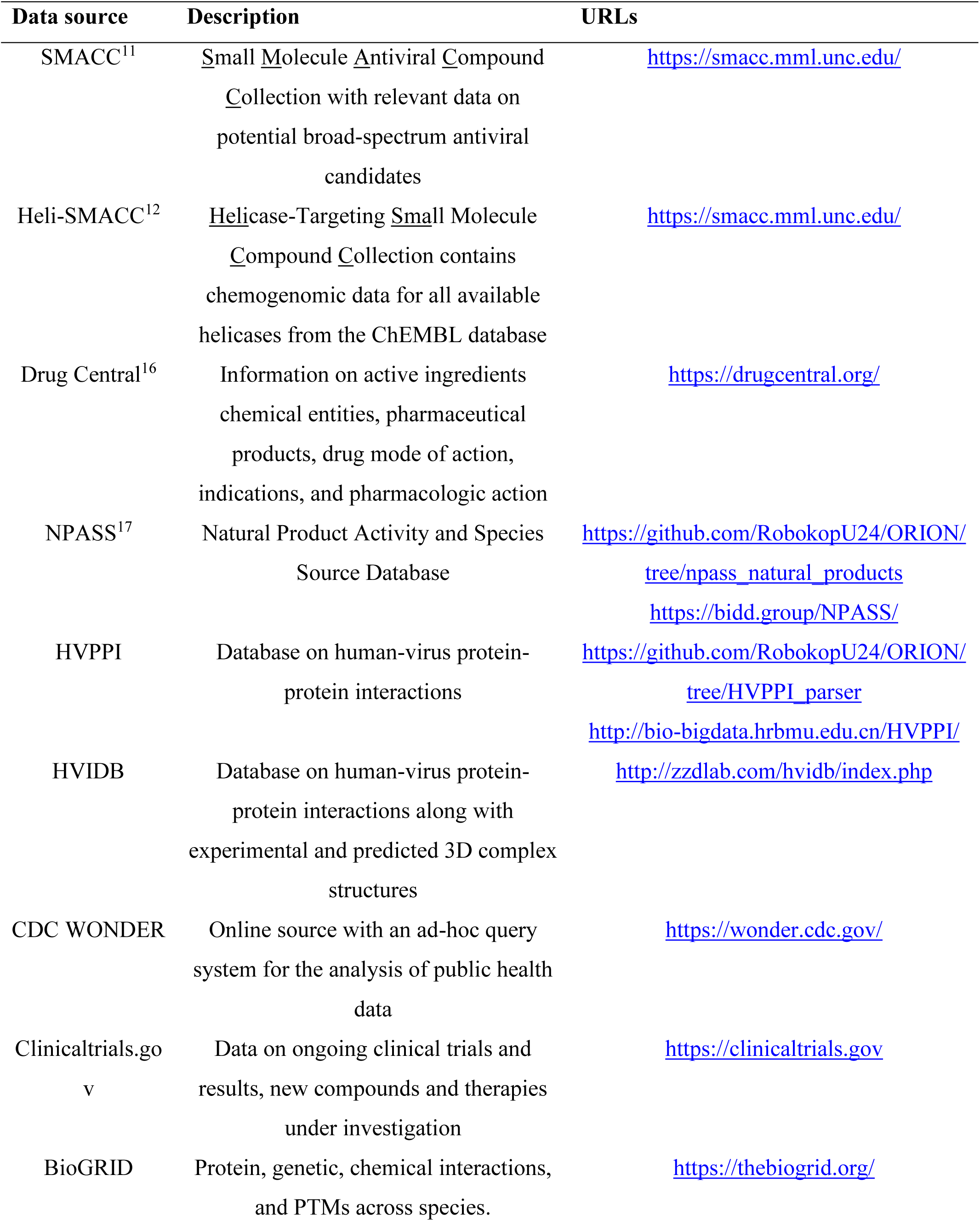
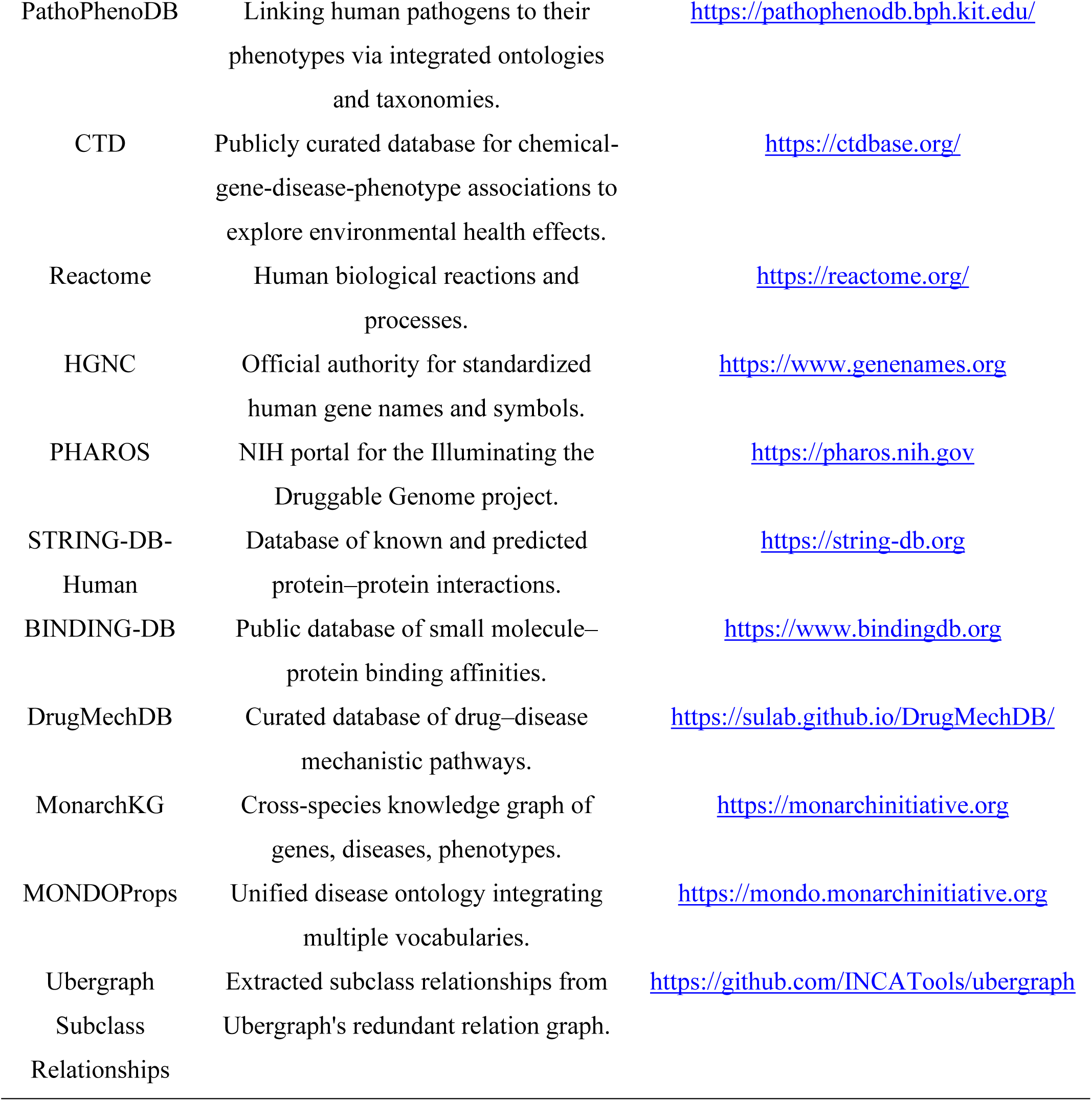
Data sources for collecting, curating, and integrating relevant data into IDKG.

**Table 2.** Node type distribution in the knowledge graph.

| **Node Type** | **Node Count** |
|---|---|
| Organism Taxon | 2,627,442 |
| Small Molecule | 1,382,871 |
| Protein | 320,273 |
| Molecular Activity | 95,353 |
| Disease | 52,113 |
| Phenotypic Feature | 38,331 |
| Gene | 37,171 |
| Chemical Entity | 26,509 |
| Molecular Mixture | 23,737 |
| Pathway | 21,811 |
| Biological Process | 18,230 |
| Gross Anatomical Structure | 10,471 |
| Anatomical Entity | 7,152 |
| Cellular Component | 5,361 |
| Named Thing | 3,734 |
| Cell | 3,388 |
| Drug | 2,511 |
| Gene Family | 1,800 |
| Chemical Mixture | 504 |
| Nucleic Acid Entity | 284 |
| Activity And Behavior | 119 |
| Geographic Location | 62 |
| Complex Molecular Mixture | 54 |
| Other types | 40 |

Each node and edge in the graph is tagged with standardized identifiers (e.g., MeSH, UniProt, SMILES, MONDO) and annotated with provenance and source metadata. The graph is stored in Neo4j and supports Cypher querying and incremental updates. Its architecture uses standardized identifiers, provenance tracking, and Biolink-compatible semantics in accordance with FAIR-oriented design principles. Relative to ROBOKOP and COVID-KOP, IDKG adds infectious-disease-focused datasets and curation workflows spanning antiviral and antibacterial activity, host– pathogen interactions, clinical information, and literature-derived evidence.

### Data Curation: SMACC, Heli-SMACC, and Chemical Mixtures

High-quality data curation was foundational to the development of IDKG. For the SMACC database, which includes both target-based and phenotypic antiviral assays, all entries with ambiguous outcomes (e.g., "inconclusive," "unclear") were removed. Post-curation, 11,108 target-based and 20,781 cell-based assay records were retained, representing 99% and 97% of the original datasets, respectively. The Heli-SMACC database, which focuses on helicase-targeting small molecules, retained 13,464 curated entries (66% of the original).

A specialized workflow was also developed to handle complex mixtures and formulations. This workflow normalized component concentrations, removed salts and undefined structures, and addressed duplicates by evaluating compositional and proportional consistency. Mixtures with identical ingredients in different proportions were retained as distinct entities. Several steps were semi-automated and supplemented with manual validation to ensure accuracy. These rigorous curation steps significantly enhance the reliability of downstream analytics. Structural normalization and duplicate handling are particularly critical for the utility of IDKG in cheminformatics and virtual screening applications. The ability to handle mixtures and biologics further expands the platform’s applicability in real-world therapeutic contexts.

### Case Study: COVID-19 and Remdesivir

To demonstrate the practical utility of IDKG for mechanistic exploration and drug repurposing, we examined remdesivir in the context of COVID-19 (Figure 2). Remdesivir was selected as a well-characterized antiviral for which molecular and clinical evidence could be compared with graph-derived connections. The example therefore serves as a proof-of-concept for connecting molecular targets, biological processes, phenotypes, and disease outcomes using COPs. The Figure 2 query can be reproduced using the example provided in the IDKG GitHub repository: https://github.com/molecularmodelinglab/ORION_IDKG/blob/master/README-IDKG.md.

**Figure 2:**
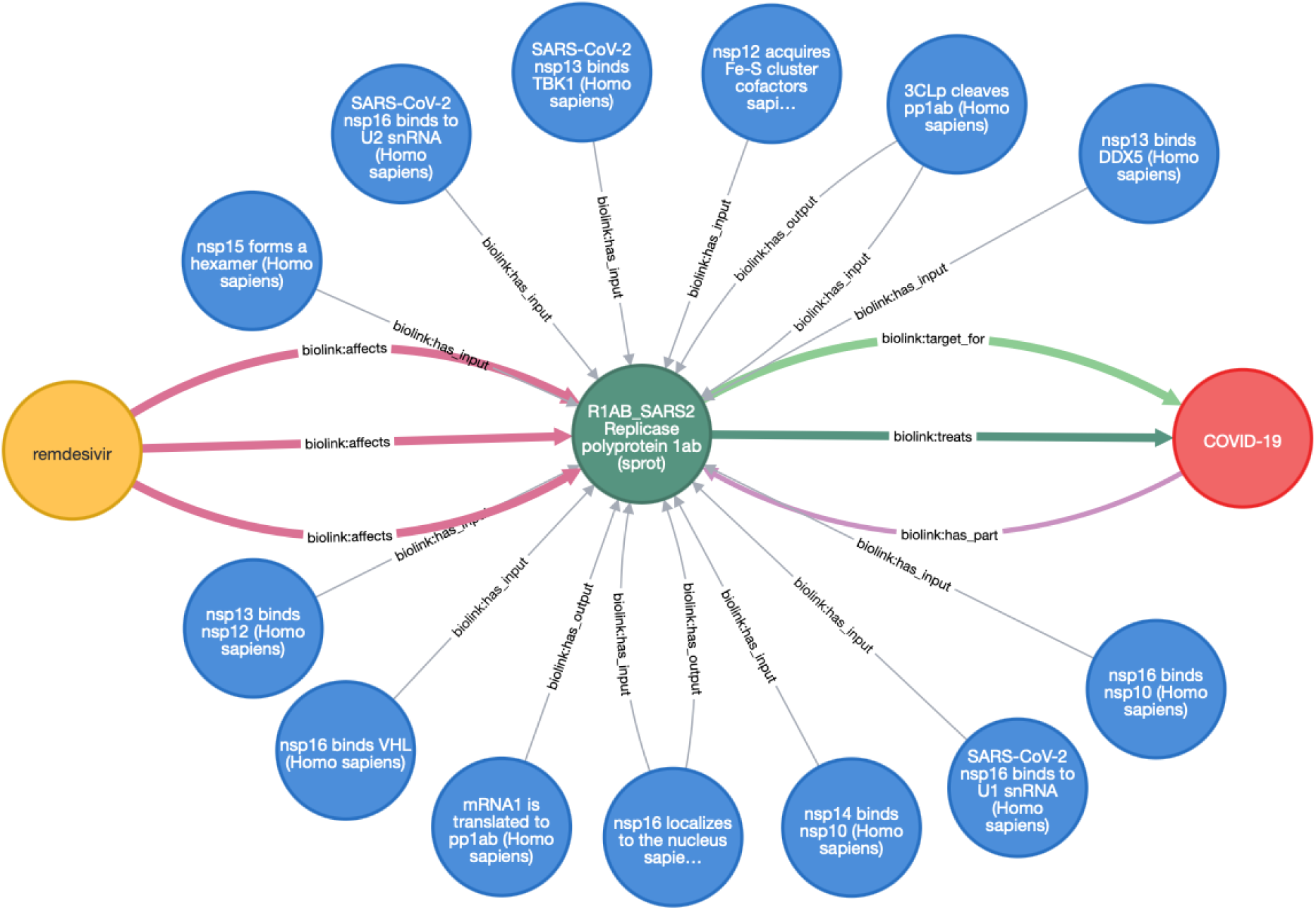
Example Clinical Outcome Pathway (COP) connecting remdesivir, SARS-CoV-2 replication-related entities, and COVID-19 phenotypes in IDKG. The displayed network is a graph-derived, literature-supported representation intended for mechanistic exploration; individual edges should be interpreted according to their source and provenance rather than as a single experimentally established causal chain.

Within IDKG, remdesivir is connected to the SARS-CoV-2 replicase polyprotein 1ab (UniProt ID: P0DTD1), from which nonstructural proteins including the RNA-dependent RNA polymerase (nsp12/RdRp) are processed. The graph representation should therefore be interpreted as an ontology- and identifier-mediated connection to the viral replication machinery rather than as a claim that every displayed edge represents a directly measured drug–protein interaction. Using COP mining, we traced a multi-step path from antiviral target-level information toward phenotypic outcomes relevant to COVID-19. The path included reduced viral replication and downstream inflammatory and respiratory phenotypes. Supporting literature for individual or adjacent relationships was identified with Chemotext and Abstract Sifter; more than 25 PubMed-indexed studies supported components of the illustrated path. Literature co-occurrence was used as supporting evidence and prioritization information. Source identifiers and provenance were retained to support traceability of the underlying assertions.

The remdesivir COP recapitulated established antiviral biology and identified additional host-response relationships associated in the graph with viral replication and disease severity, including NF-κB signaling, pro-inflammatory cytokines such as IL-6 and TNF-α, and pulmonary epithelial phenotypes. These relationships are best interpreted as literature-supported hypotheses connecting viral inhibition to host-response outcomes rather than as experimentally validated downstream effects of remdesivir within this study. The path also highlights measurable clinical or translational variables, including viral load, inflammatory markers, and oxygenation indices, that may be useful when evaluating related mechanistic hypotheses.

The remdesivir example therefore provides a proof-of-concept for generating mechanistically interpretable, literature-supported paths that connect molecular interactions with clinical phenotypes. Similar COPs can be generated for other antiviral agents and compared with remdesivir to identify shared or divergent mechanistic features. Such comparisons are intended to prioritize hypotheses for subsequent experimental or clinical evaluation rather than to establish efficacy or causal mechanism directly from graph structure.

### Case Study: QSIs and Synergistic Antibiotic Pairs

To illustrate the Infectious Disease Knowledge Graph (IDKG)’s potential to support translational research and antimicrobial innovation, we conducted an integrated case study on quorum sensing inhibition (QSI) in multidrug-resistant (MDR) bacterial pathogens, specifically Pseudomonas aeruginosa and Acinetobacter baumannii (Figure 3). Quorum sensing (QS) is a bacterial cell–cell communication process critical for virulence, biofilm formation, and resistance phenotype expression in many Gram-negative organisms. Disrupting QS pathways offers a promising non-bactericidal strategy to enhance the efficacy of conventional antibiotics.

**Figure 3:**
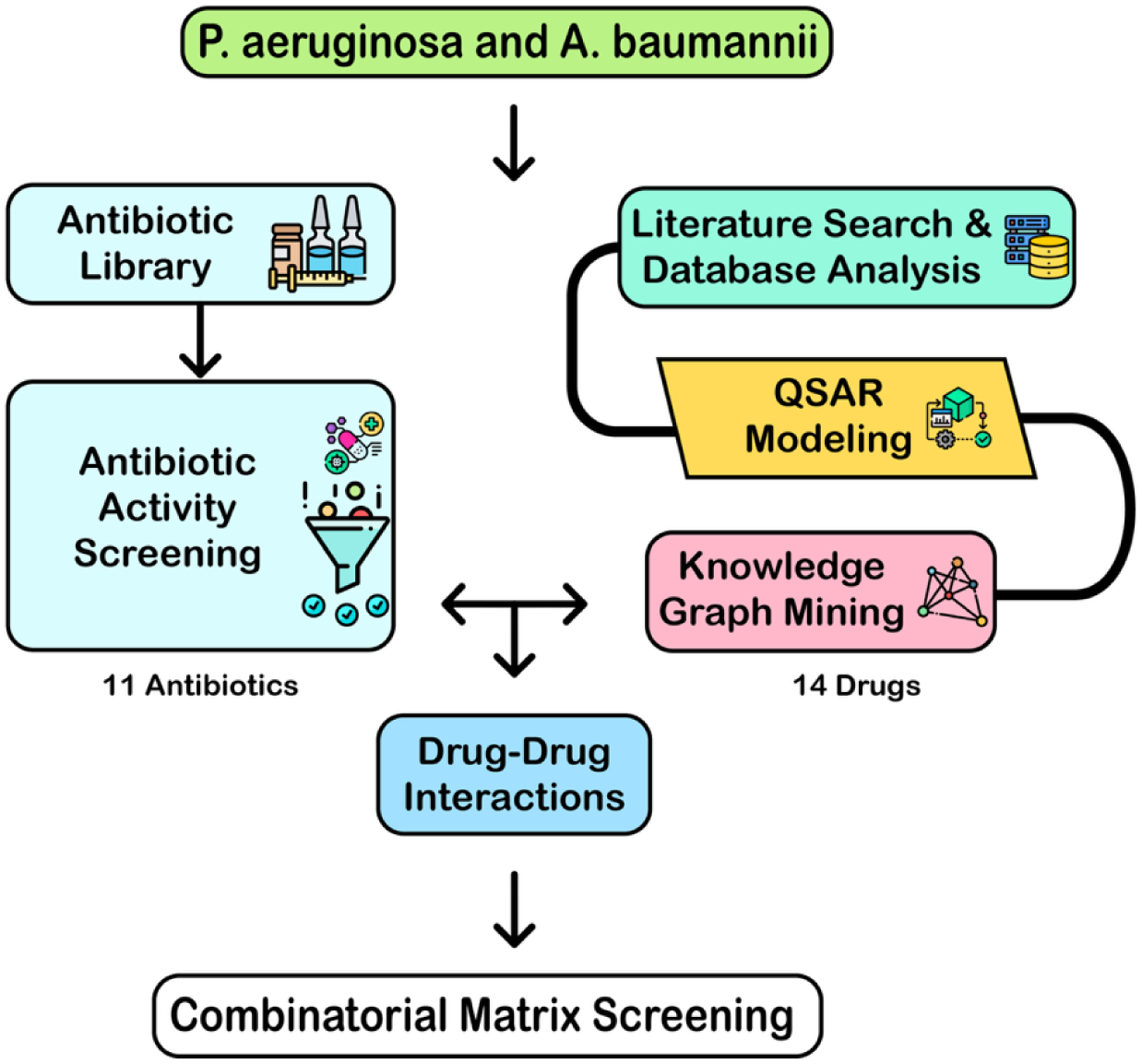
Workflow for the quorum-sensing inhibition case study. Literature and database mining informed QSAR modeling and candidate selection; selected compounds were evaluated with antibiotics by combinatorial matrix screening, and IDKG was subsequently used to generate mechanistic hypotheses for experimentally observed interactions.

The quorum-sensing inhibitor (QSI) modeling and screening workflow has been reported in detail previously. ChEMBL was searched for assays containing "quorum sensing"; relevant P. aeruginosa data were curated in KNIME v4.5.2, while no suitable A. baumannii assay data were identified for QSAR modeling. Entries were classified as active when inhibition, activity, or efficacy was at least 50%, or when ChEMBL annotations indicated an active or dose-dependent effect. After removal of salts, mixtures, duplicates, and structure standardization, the final training set contained 372 compounds (36 active and 336 inactive). A Random Forest QSAR model was evaluated using five-fold external cross-validation, yielding an AUC-ROC of 0.89; the corresponding model trained after randomization of activity labels yielded an AUC-ROC of 0.51. The validated model was then applied to 12,584 approved and investigational compounds in the Inxight pharmaceutical collection, of which 49 were predicted active, and candidates were further prioritized using drug–drug interaction predictions and knowledge-graph/literature evidence.^21^

Selected known and predicted QSIs were tested with 11 antibiotics against five P. aeruginosa and five A. baumannii strains. The combination assay used 1536-well plates with six-point titrations and final concentrations from 46 to 0.18 μM; bacterial growth was quantified by OD600. The prior study evaluated combination response profiles using DBSumNeg, defined as the summed difference between observed and expected growth-inhibition responses, with DBSumNeg < −3 used as the synergy criterion. Review of the current version of the study identified 26 combinations meeting this threshold in at least one tested strain; 22 combinations showed synergy in at least one strain from each bacterial species. Piperacillin combinations with piroxicam, indomethacin, and ketoprofen showed synergy in both P. aeruginosa and A. baumannii strains. Full response profiles and tabulated synergistic and antagonistic interactions are available in the associated supplemental data.^21^

To explore candidate mechanisms underlying the observed synergies, we used IDKG to construct COPs linking QSIs and antibiotics to phenotypic outcomes (Figure 5). The resulting paths implicated quorum-sensing regulators (e.g., LasR and RhlR), multidrug efflux systems (e.g., MexAB-OprM), membrane-permeability processes, and virulence-associated pathways as plausible mediators of antibiotic sensitization. Literature mining with Chemotext and Abstract Sifter identified co-occurrence support for candidate compounds and these pathways. These graph-derived relationships should be interpreted as mechanistic hypotheses requiring direct experimental validation; the present study does not establish that curcumin, naldemedine, or the other tested compounds act through any single inferred target or pathway.

**Figure 4:**
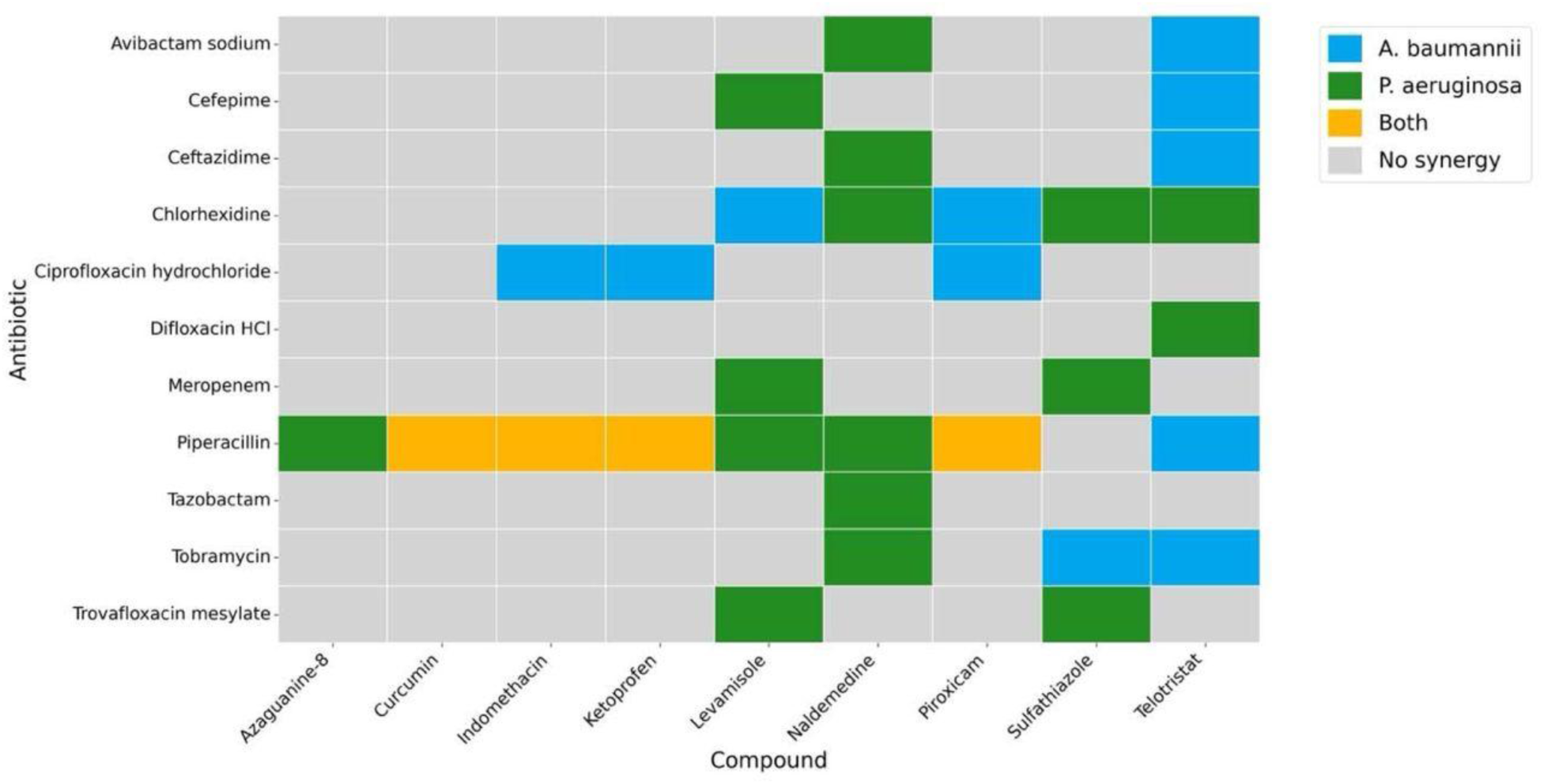
Synergistic interactions between antibiotics and candidate compounds across A. baumannii and P. aeruginosa strains. Cells summarize combinations meeting the predefined study synergy criterion (DBSumNeg < −3); colors indicate the organism group in which synergy was observed, as defined in the figure key.

**Figure 5:**
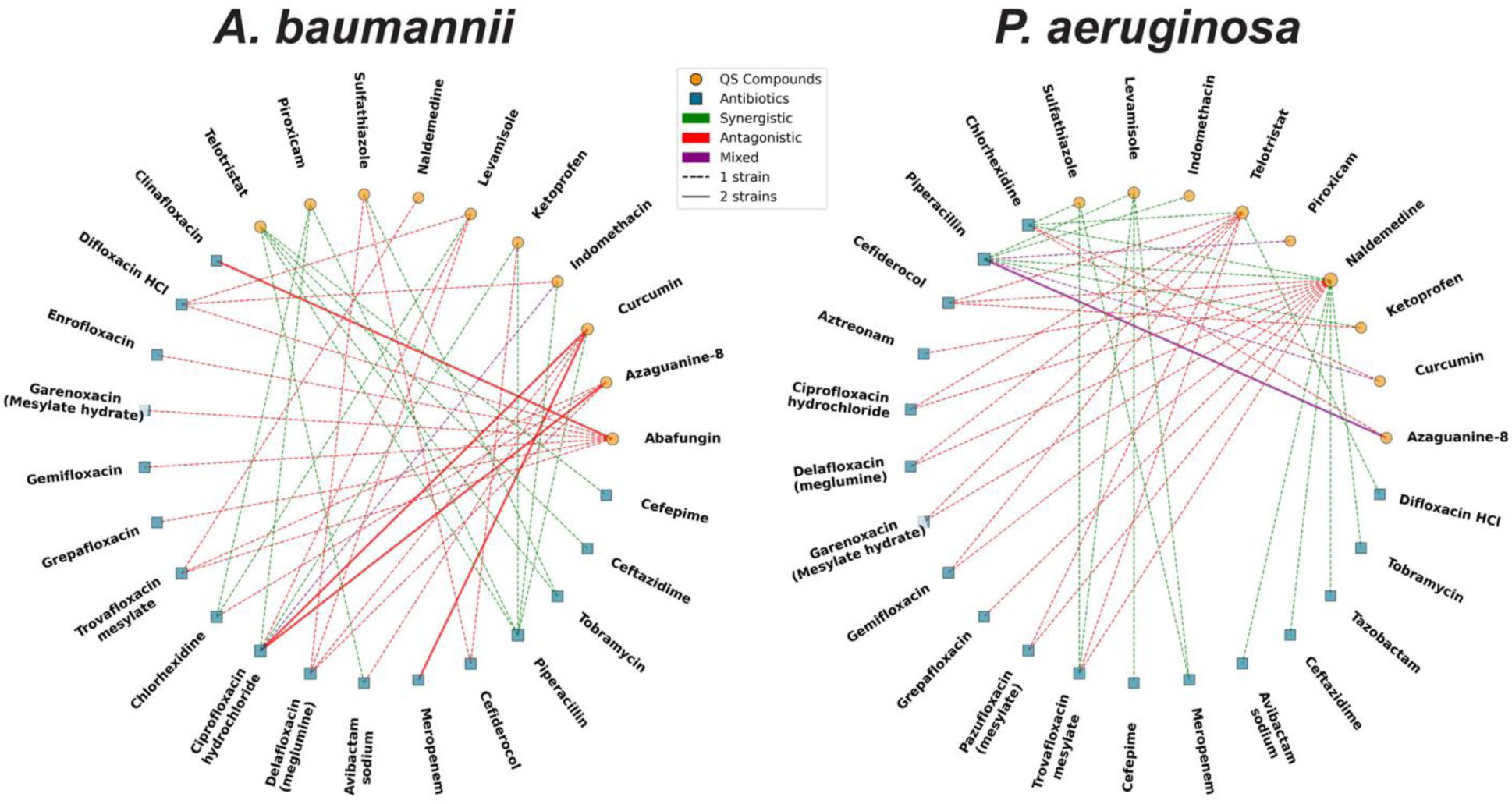
Interaction networks for A. baumannii (left) and P. aeruginosa (right) integrating antibiotics, candidate quorum-sensing inhibitors, and experimentally classified combination outcomes. Green and red edges denote synergistic and antagonistic interactions, respectively; graph-derived mechanistic relationships are interpreted as hypotheses for follow-up validation.

This case study demonstrates how IDKG can connect cheminformatics modeling, experimental screening results, and graph-based reasoning to support interpretation of synergistic antimicrobial combinations. The observed synergy is established by the experimental matrix screening, whereas IDKG provides candidate mechanistic explanations that can guide follow-up studies. These hypotheses provide a basis for direct target-level, pathway-level, and in vivo validation of selected combinations.

### Platform Usability

To maximize accessibility and usability, the Infectious Disease Knowledge Graph (IDKG, https://idkg.mml.unc.edu) was developed as an interactive, web-based platform designed for researchers, clinicians, and data scientists. The graph was deployed within a Neo4j database environment, enabling efficient execution of complex Cypher queries. A user-friendly graphical interface was implemented to allow intuitive exploration of drug–target–disease relationships, clinical outcome pathways, and compound synergy networks without requiring programming expertise. The interface includes a query construction tool that builds Cypher queries via user-fillable fields and dropdown boxes, enabling users to design up to 10 query paths, each with a start node, end node, and up to five intermediate nodes, specifying node names, identifiers, and predicates for flexible, detailed graph exploration. A node search function assists users in finding node names and IDs within the graph by providing suggestions based on partial or full name and identifier input, which can be directly copied into queries for streamlined construction. Query results appear in a downloadable table where each row represents a unique answer subgraph, with options to toggle columns, view metadata by hovering, and visualize selected or multiple subgraphs to build larger networks for hypothesis exploration. Additionally, modular data views were created to support filtering by pathogen, therapeutic class, assay type, or evidence level. The platform architecture supported interoperability with cheminformatics and systems biology tools, and data export is provided in CSV format to facilitate downstream analysis. Together, these features support practical use of IDKG across diverse research and translational settings.

## Experimental

### Data Collection and Curation

To build the IDKG, we collected data from 20 biomedical repositories relevant to infectious diseases (Table 1), including SMACC, Heli-SMACC, DrugCentral, HVPPI, HVIDB, NPASS, ClinicalTrials.gov, and CDC resources. Entities were mapped using standardized identifiers and curated using open-source pipelines with an emphasis on quality control, reproducibility, and chemical-structure integrity. Data integration and normalization were performed using the Operational Routine for the Ingest and Output of Networks (ORION) pipeline (https://github.com/RobokopU24/ORION), building on the graph-integration framework used for ROBOKOP.8 We developed a dedicated fork (https://github.com/molecularmodelinglab/ORION_IDKG), with included data sources and graph-creation parameters defined in the idkg-graph-spec.yml file. New parsers were implemented for SMACC, Heli-SMACC, HVIDB, NPASS, BioGRID, PathoPhenoDB, CDC WONDER, and HVPPI. Integration of each new data source involved five steps: (1) source-structure analysis; (2) filtering and missing-value handling; (3) identifier standardization using common biomedical ontologies; (4) entity classification; and (5) mapping of source-specific relationships to Biolink Model predicates. SMACC and Heli-SMACC curation followed previously published workflows.11,12 Additional curation code is available at https://github.com/cleber-mf/data_curation. The IDKG architecture was designed in accordance with FAIR principles^20^ through standardized transformations, provenance tracking, modular processing, and machine-readable identifiers; this statement describes design intent and implemented practices rather than a formal claim of complete FAIR compliance.

In addition to individual compound records, we developed and implemented a specialized workflow for handling mixtures and drug formulations. These datasets typically list active ingredients and their proportions but pose unique challenges for normalization and deduplication. The workflow includes automated filtering of entries with incomplete metadata (e.g., missing concentrations or endpoint values), removal of salts, oils, macromolecules, and undefined chemical structures, as well as normalization of chemotypes to ensure structural consistency across the dataset. Duplicate analysis for mixtures was adapted to recognize that identical constituents in different proportions are not replicates; only formulations with exact compositional and proportional matches are flagged as duplicates. Several steps of the mixture curation process, particularly duplicate detection, involve semi-automated protocols with manual verification to ensure data quality. Ongoing improvements aim to further automate these tasks. The full mixtures curation pipeline is also publicly available at https://github.com/cleber-mf/data_curation.

### Data Integration and Graph Construction

Data integration for IDKG followed a systematic workflow designed to harmonize curated records from heterogeneous biomedical repositories into a unified, queryable structure. After collection and curation, entities including chemicals, proteins, phenotypes, and diseases were assigned standardized identifiers (e.g., SMILES, UniProt, MeSH, and MONDO, as appropriate to entity type), and relationship assertions were mapped into a centralized Assertion Table (Table 3). Each assertion retained its source, entity identifiers, relationship type, and relationship-specific details such as “treats” or “causes.” Literature-derived associations from Chemotext and Abstract Sifter were incorporated as co-occurrence-based evidence and were kept conceptually distinct from experimentally established causal relationships. The integrated graph contains more than 4.6 million nodes and 26 million edges in Neo4j. Node types were assigned using the most specific applicable Biolink Model category according to the class hierarchy. Relative to the baseline ROBOKOP graph, IDKG adds infectious-disease-focused datasets and curation workflows supporting graph mining, pathway exploration, and hypothesis generation.

**Table 3.**
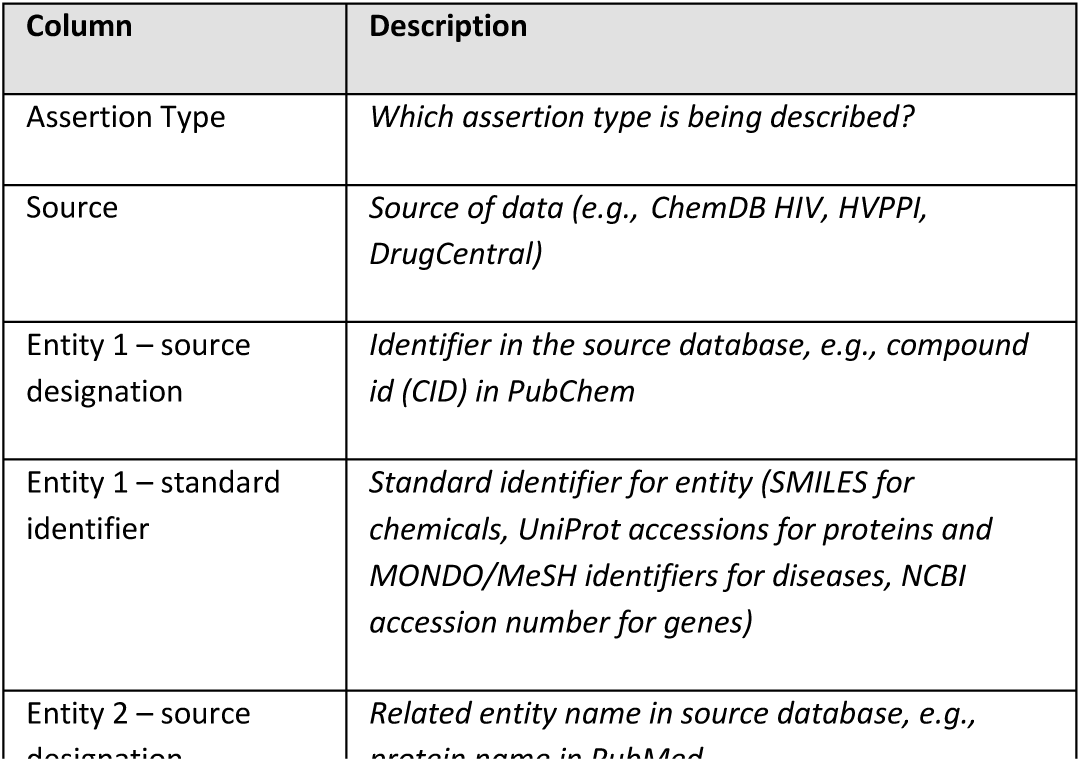
Assertion Table Layout.

| Column | Description |
|---|---|
| Assertion Type | *Which assertion type is being described?* |
| Source | *Source of data (e.g., ChemDB HIV, HVPPI, DrugCentral)* |
| Entity 1 – source designation | *Identifier in the source database, e.g., compound id (CID) in PubChem* |
| Entity 1 – standard identifier | *Standard identifier for entity (SMILES for chemicals, UniProt accessions for proteins and MONDO/MeSH identifiers for diseases, NCBI accession number for genes)* |
| Entity 2 – source designation | *Related entity name in source database, e.g., protein name in PubMed* |

### Knowledge Graph Mining and Reasoning

To extract actionable insights from the Infectious Disease Knowledge Graph (IDKG), we implemented a suite of complementary reasoning and mining tools designed to support target identification, drug repurposing, and mechanistic hypothesis generation. These tools enable both rule-based and machine learning–driven exploration of relationships among drugs, targets, pathways, phenotypes, and infectious disease outcomes.

### 1. Clinical Outcome Pathways (COPs)

One of the graph-based reasoning approaches implemented in IDKG is the construction of Clinical Outcome Pathways (COPs). COPs were previously formalized as a sequence linking a molecular initiating event through one or more intermediate molecular or cellular events to a clinical outcome, and ROBOKOP was used to demonstrate their extraction from biomedical knowledge graphs.^22^ In IDKG, candidate mechanistic paths are generated using Cypher queries in Neo4j. Path exploration follows the ExEmPLAR framework, which supports construction of multi-step knowledge-graph queries, visualization and export of answer subgraphs, and ranking of candidate paths using PubMed abstract co-mention counts for selected node pairs or triplets.^23^ These literature-based rankings are used as evidence-prioritization heuristics and should not be interpreted as calibrated estimates of causal probability.

### 2. Literature-Augmented Validation

Candidate paths and hypotheses were evaluated using literature co-occurrence analysis. Chemotext and Abstract Sifter were used to identify publications that co-mentioned candidate entities, while the ExEmPLAR workflow provides PubMed co-mention counts that can be used to sort and prioritize answer paths.^23^ This procedure provides a semi-automated evidence-prioritization layer; co-occurrence alone does not establish directionality, causality, or experimental validation of an inferred pathway.

## Limitations

IDKG inherits limitations from its underlying data sources, including differences in update frequency, evidence quality, ontology coverage, experimental context, and reporting conventions. Identifier normalization and semantic mapping can introduce uncertainty, particularly for pathogen strains, complex mixtures, and entities represented differently across databases. Literature co-occurrence provides supporting evidence for graph-derived relationships but should not be interpreted as proof of causality, and COPs are intended primarily for hypothesis generation unless their component relationships have been independently established. Coverage is not uniform across pathogens or therapeutic areas, and the graph can become temporally stale as source databases change. The QSI case study relies on a previously reported Random Forest model trained on a relatively small and imbalanced dataset (36 active and 336 inactive compounds), although five-fold external cross-validation and a randomized-label control were reported.^21^ Independent or scaffold-aware external validation would further strengthen assessment of model generalizability. Likewise, the mechanistic explanations proposed for synergistic antibacterial combinations require direct target- and pathway-level experimental validation.

## Conclusions

IDKG integrates 20 biomedical, chemical, host–pathogen, and clinical data sources into an infectious-disease-focused knowledge graph containing more than 4.6 million nodes and 26 million edges. Curation workflows for small molecules, biologics, and mixtures, together with Biolink-compatible semantic mapping and provenance-aware integration, support graph-based exploration of drug–target–disease relationships. The remdesivir and antibacterial case studies illustrate complementary uses of the resource: reconstruction of literature-supported mechanistic paths and interpretation of experimentally observed drug-combination phenotypes. These examples support IDKG as a framework for hypothesis generation and prioritization rather than as a substitute for experimental causal validation. The web interface and Neo4j query environment provide multiple routes for interacting with the graph, and the system was designed in accordance with FAIR principles. Future development will focus on broader pathogen coverage, deeper clinical annotation, improved graph versioning and validation, ADMETox integration, de novo design, and more explicit causal-reasoning methods.

## Author contributions

H.J.M., J.M.B., and V.K. contributed to data curation, formal analysis, methodology, software development, and visualization. V.K. additionally led knowledge graph construction and deployment. H.J.M. and V.K. contributed to writing the original draft. R.S.T., H.J.M, E.M., and A.T. contributed to manuscript review and editing. E.M., K.C., and A.T. were responsible for conceptualization, funding acquisition, and supervision.

## Data availability

The IDKG web platform is available at https://idkg.mml.unc.edu. Source code and graph-construction specifications for the IDKG implementation of ORION are available at https://github.com/molecularmodelinglab/ORION_IDKG, and the curation workflows used for SMACC, Heli-SMACC, and related chemical data are available at https://github.com/cleber-mf/data_curation. Data incorporated from third-party repositories remain subject to the access terms and licenses of the originating resources. The manuscript describes the current graph build and provides the information required to reproduce the included example queries; additional project-specific materials may be obtained from the corresponding authors where redistribution rights permit.

## Acknowledgements

This work was supported by the National Institute of Allergy and Infectious Diseases (NIAID), NIH/HHS, under award 75N93024C00038; and by the National Institute of Environmental Health Sciences (NIEHS), NIH/HHS, under awards R41 ES033857 and U24 ES035214.

